# Frontal Midline Theta Reflects Cognitive Control During Planning

**DOI:** 10.1101/648758

**Authors:** Marcos Domic-Siede, Martín Irani, Joaquín Valdés, Marcela Perrone-Bertolotti, Tomás Ossandón

**Affiliations:** Neurodynamics of Cognition Laboratory, Departamento de Psiquiatría, Pontificia Universidad Católica de Chile, 8320000 Santiago, Chile; Laboratory for Brain-Machine Interfaces and Neuromodulation, Departamento de Psiquiatría, Pontificia Universidad Católica de Chile, 8320000 Santiago, Chile; Univ. Grenoble Alpes, Univ. Savoie Mont Blanc, CNRS, LPNC, 38000 Grenoble, France

**Keywords:** planning, frontal midline theta frequency band (FMθ), time-frequency analysis, prefrontal cortex, cognitive control

## Abstract

Neural correlates of cognitive planning are not understood well at present. Behavioral paradigms targeting this function are a current challenge in cognitive neuroscience. We recorded EEG activity while subjects were performing a novel behavioral paradigm that evaluates cognitive planning function. Participants showed longer reaction times and decreased accuracy during the planning condition compared to the control condition, suggesting that the planning condition is more time consuming, therefore reflecting higher cognitive demand. Moreover, cognitive planning induced a frontal midline theta (FMθ) frequency band originating in prefrontal cortex (PFC) as shown in previous cognitive control studies. When subjects began planning, there was a progressive and sustained increase in FMθ starting after 500 milliseconds (ms) of planning. Hence, we characterized for first time, both spatial and temporal FMθ dynamics of cognitive planning as a marker of cognitive control function.

## 1. Introduction

Cognitive control or Executive functions (EFs) are a theoretical construct that include a wide range of higher order cognitive functions associated with goal-directed behavior (Lezak, 1995; Shallice, 1991; Stuss, 1992; Zwosta, Ruge & Wolfensteller, 2015; Cooper, 2010). One of the high-order cognitive control function is planning (Collins & Koechlin, 2012; Sira & Mateer, 2014; Lunt et al. 2012), which consists on the ability of developing a sequenced plan to achieve a goal in an organized, strategic and efficient manner (Hayes-Roth & Hayes-Roth, 1979). Planning allows imagining what the future might be and how our behavior could affect and change the current state leading us to this imagined future (Benson, 1993). The extent of plans can range from a simple motor behavior (e.g. planning a sequence of key presses) (Pascual-Leone et al., 1993) to a high-demanding cognitive task (e.g. deciding on the steps required to land an airplane) (Suchman, 1987). Planning behavior can be divided into two major phases: *i)* a mental planning phase that involves the internal representation of a sequence of steps (plans) (Wilensky, 1983) and *ii)* a planning execution phase that involves the motor action to achieve a goal previously planned (Grafman & Hendler, 1991). Thus, planning can be measured in simple and/or more complex tasks (Schwartz et al., 1991). Typically, in the context of higher order cognitive processes, planning requires the operation of several components of the EFs (e.g., working memory, attentional control, response inhibition) making the experimental manipulation and isolated measurement of other EFs difficult (Hayes-Roth & Hayes-Roth, 1979; Tremblay et al, 1994).

Since planning is compromised in several psychiatric and cognitive disorders such as Attention-Deficit Hyperactivity Disorder (Barkley, 2004; Gau & Shang, 2010), Major Depressive Disorder (Bora, Harrison, Yücel & Pantelis, 2013; Rive, Koeter, Veltman, Schene & Ruhé, 2016), Bipolar Disorder (Rive et al., 2016), Schizophrenia (Holt, Wolf, Funke, Weisbrod & Kaiser, 2013), frontotemporal dementias (Lima-Silva et al., 2013) as well as in frontal lesions (Karnath, Wallesch & Zimmermann, 1991), the implementation of proper experimental tests has been challenging for clinical neuropsychology which has contributed considerably to the study of planning, especially in the design of behavioral paradigms which allow quantifying and characterizing normal and impaired planning performance in healthy and pathological subjects.

Neuroimaging studies have provided valuable evidence about the critical role of the prefrontal cortex (PFC) in cognitive control, including planning. The most traditional paradigm used in those studies to evaluate planning is the Tower of London Task (TOL) (Shallice, 1982; Unterrainer et al., 2004) which suggested an important implication of the dorsolateral PFC (DLPFC) (Nitschke et al., 2017), the dorsal anterior cingulate cortex (dACC) and the superior parietal lobe (SPL), among other brain regions (Kirsch et al., 2006; Newman, Carpenter, Varma, & Just, 2003; Owen, Doyon, Petrides & Evans, 1996). Besides TOL, there are others traditional tests that measure planning: Porteus (1959) proposed the Porteus Maze Task which has been widely used to study planning skills in a PFC-dependent visuospatial context in healthy control and neuropsychiatric population (Gallhofer, Bauer, Lis, Krieger, & Gruppe, 1996; Krieger, Lis, & Gallhofer, 2001; Lee, Chou, Li, Wan, & Yen, 2007; Lezak, 1995; Peters & Jones, 1951; Tremblay et al., 1994). The tasks mentioned above have a major problem of reduction of the ecological validity (i.e. proposing natural contexts or activities well-connected with everyday life situations), because in order to attempt to control confounding factors, paradigms become more artificial and may have lesser predictive validity (Miotto & Morris, 1998; Burgess, Simons, Coates & Channon, 2005; Oosterman, Wijers, & Kessels, 2013; Campbell el al., 2009). To address this problem, more ecological tasks analogous to real-world planning situations (Miotto & Morris, 1998; Burgess, Simons, Coates & Channon, 2005) have been proposed. Noticeably, Wilson et al. designed the Behavioral Assessment of the Dysexecutive Syndrome battery (BADS) (Wilson, Alderman, Burgess, Emslie & Evans, 1996) to measure EFs including a subtest called Zoo Map Task that provides a valid planning ability indicator (Oosterman, Wijers, & Kessels, 2013). Importantly, this subtest has the advantage of providing a planning and organizational skills measurement in a more ecological way. In the present study, in order to evaluate cognitive planning function, we used an adaptation of Porteus Maze and Zoo Map Task paradigm.

While the precise brain regions involved during planning are amenable to imaging studies that use fMRI or PET, its fine temporal and neural properties remain elusive. In this study, we address this issue by analyzing neuronal oscillatory activity. We hypothesize that FMθ could be a physiological mechanism of temporal dynamics reflecting cognitive planning processes (Buzsáki & Draguhn, 2004). Over the past 15 years, there has been a strong research focus on FMθ activity with scalp EEG assessment, which has been associated closely to several cognitive control functions such as working memory and attentional control (Cavanagh & Frank, 2014; Deiber et al., 2007; Green & McDonald, 2008; Onton, Delorme, & Makeig, 2005; Summerfield & Mangels, 2005; White, Congedo, Ciorciari, & Silberstein, 2012; Raghavachari et al., 2006). Furthermore, FMθ has been posited as a candidate mechanism by which cognitive control might be biophysically performed (Cavanagh & Frank, 2014). However, the dynamic interplay between EEG oscillatory activity and planning function remains unknown. Under this context, the present study attempts to answer whether cognitive planning implementation induces a FMθ band activity originated in PFC using a novel and ecological experimental paradigm.

## 2. Materials and Methods

### 2.1. Participants

Data was collected from twenty-seven right-handed healthy adults (13 females) between 19 to 38 years-old (mean age = 27.81, standard deviation (SD) = 4.58 years). The sample size was calculated using G*Power 3.1.9.2 software (http://www.gpower.hhu.de/) considering statistical Wilcoxon match-paired test, effect size of 0.7, alpha value of 0.05, and a power of 0.95 (Faul, Erdfelder, Lang & Buchner, 2007). No participant reported neurological or psychiatric disorders according to the International Neuropsychiatric Interview, Spanish version adapted (Ferrando, Bobes, Gibert, & Soto, 2000). All participants had normal or corrected-to-normal vision. They were paid CLP$10,000 (approximately USD$15.76 or €13.30) for their participation. Procedures were approved by the bioethics committee of the Faculty of Medicine of Pontificia Universidad Católica de Chile and all participants signed an informed consent form prior to the beginning of the study (research project number: 16-251).

### 2.2. Experimental Design and Procedure

We created a planning task paradigm based on Zoo Map Task (Wilson et al., 1996) and Porteus Maze (Porteus, 1959) programmed in the Presentation Software® by Neurobehavioral Systems (Version 18.0, www.neurobs.com, Neurobehavioral Systems, Inc., Albany, CA) and stimuli were designed using open source SVG tool Inkscape (www.inkscape.org). The task was coupled to an EEG scalp system and an eye-tracker (ET) system allowing to give an online feedback of eye movements. The experiment is composed of two conditions: a planning condition and a control condition, each of which included three different periods (see below). These conditions were constructed with a similar structure that allowed control of confounding factors and perceptive components involved in the task and thus, help improve the specific assessment of the processing involved in cognitive planning. Stimuli were projected on an ASUS VG248QE 24” LCD monitor located 82 cm away from the subject.

#### 2.2.1. Planning Condition

The planning condition consisted of 36 trials each with a distinct gray-scale maze that represent a zoo map, preceded by three seconds of a centred fixation cross as a baseline. Inside the maze were a gateway and several paths leading to locations of four animals (see Figure 1A). Trials were pseudo-randomized. The planning condition was composed of three different periods: planning, execution and response (see Figure 2A).

**Figure 1.**
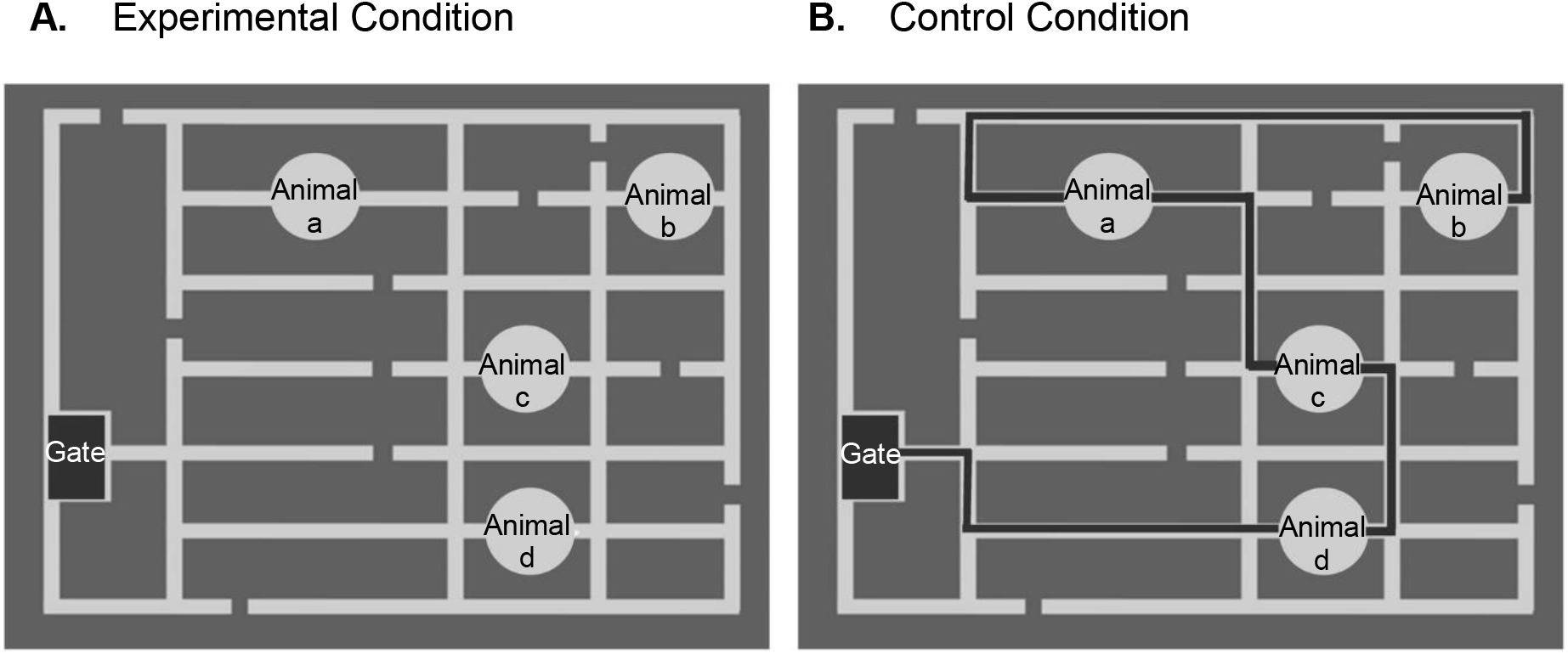
Stimuli of Planning and Control Condition. Illustrative example of the task stimuli is shown. Each stimulus consists of a zoo map with a starting gate, four animals located along the maze and different paths that may or may not lead to their locations. During the experimental condition (A) subjects had to plan a path from the gate passing through all animal locations, considering a set of rules. On the other hand, for the control condition (B) a marked line indicating an already existing path was shown (black line). Here, subjects were instructed to look at this path and figure out whether the rules were followed or not.

**Figure 2.**
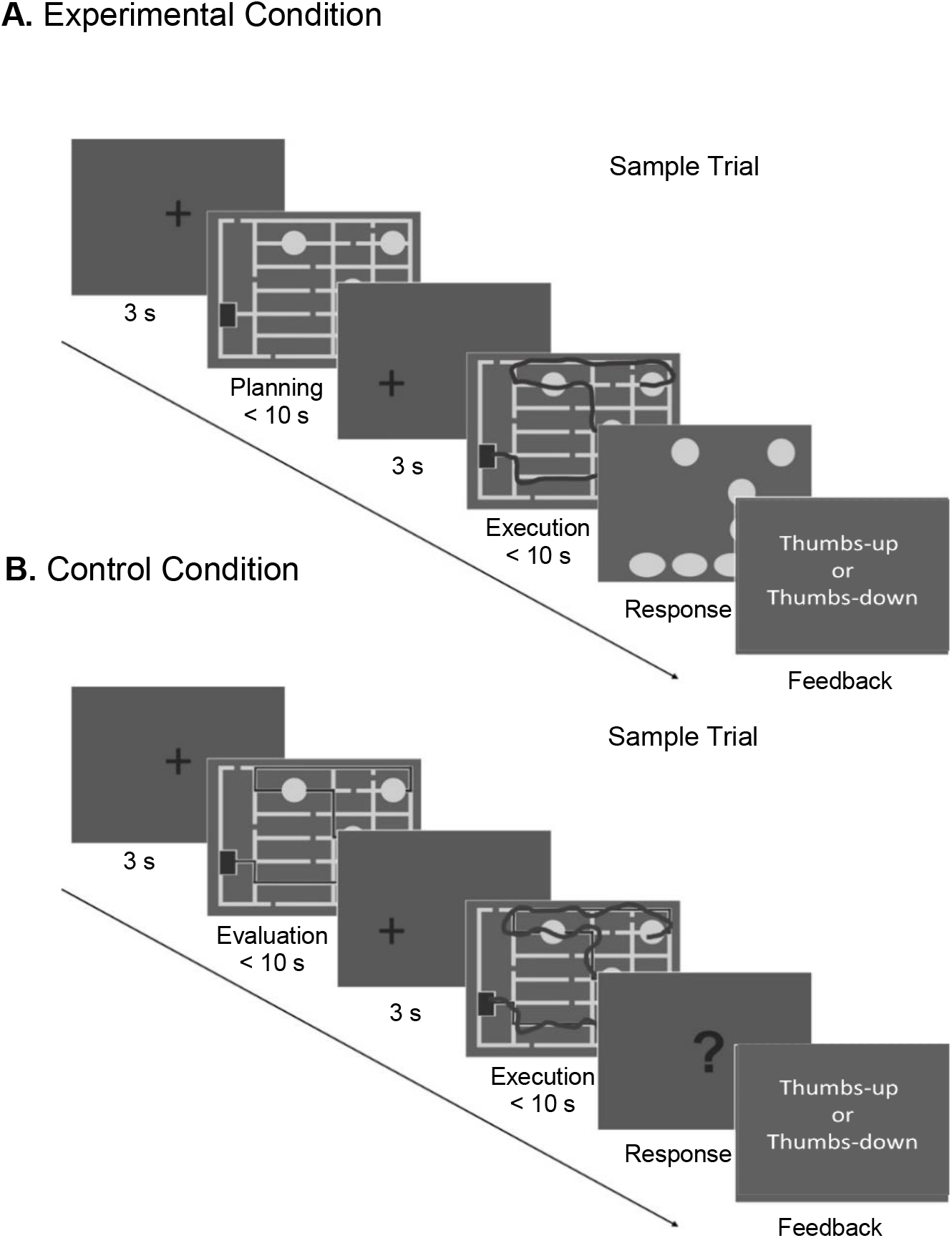
Experimental Design. Typical planning condition trial. Planning trial started with a fixation cross presented for 3 seconds. Subjects were then instructed to plan a path visiting all the four animal locations with a maximum time of 10 seconds, following a set of rules (previously explained). Afterwards, a shifted fixation cross was shown. Once the maze appeared again, subjects had to execute the trace planned in the previous planning period using their gaze with a visual feedback that delineated their gaze movement in real-time (dark line) with a maximum time of 10 seconds. Then occurred the response period where the subjects had to indicate the sequence made during execution by arranging the animals in the chosen order with a joystick. Based on their response, subjects received feedback (thumbs-up when correct and thumbs-down when incorrect). B) Typical control condition trial. A fixation cross appeared for 3 seconds. Next, subjects were instructed to look at an existing traced path (dark line) and evaluate whether it followed the rules or not. Next, a shifted fixation cross appeared again after which the maze reappeared. This time subjects had to replicate the already traced route having the same visual feedback as the execution planning period. Next came the response period where they had to answer if the traced sequence followed the previously stated rules or not by pressing a joystick button. Based on their response, subjects received feedback (thumbs-up when correct and thumbs down when incorrect.

##### Planning period

Subjects were instructed to find a path to complete a sequence of visits to all four animals (in any order) according to the following set of rules: (1) Plan the path as fast as possible within a maximum of 10 seconds, (2) Start from the beginning (the gateway) and conclude the path at the fourth animal visited, (3) Do not pass through the same path or corner twice, (4) Do not cross a dead end, (5) Do not cross a path perpendicularly. Planning period was over once the subject pressed a button from a joystick whenever they finished planning or if they exceeded the maximum time. Reaction time was recorded for further analyses.

##### Execution planning period

In this period, the maze was shown again and subjects were instructed to trace their previous planned path using their gaze through an online eye movement feedback given and registered by an ET system (EyeLink 1000 Plus, www.sr-research.com, SR Research, Mississauga, Ontario, CA). Calibrations of the ET were made at the beginning of the experiment and each five trials completed. Subjects had a maximum time of 10 seconds to trace the planned path but once they crossed the fourth animal visited, they could finalize by pressing a button. Their reaction times were saved for further analysis. This period was preceded by three seconds of a shifted fixation cross shown immediately after the planning period which indicated the start position (gate) of this execution period and facilitate the tracing.

##### Response planning period

After 10 seconds or at the press of the button by the end of the execution period, the maze disappeared and only the animals stayed on the screen in the same spatial location they appeared in previous periods. Additionally, there were four yellow circles at the bottom of the screen. Subjects were asked to insert the animals in each circle following the same order in which they visited them during the execution period. Then, subjects got feedback based on the feasibility of the traced sequence (thumbs-up or thumbs down when the answer was either correct or incorrect, respectively) and their accuracy was considered for analyses.

Consequently, the behavioral features used to measure planning performance were the reaction time during planning period (the time that subjects needed to figure out how to solve the maze following the rules) and the reaction time during the execution period (the time that subjects needed to execute the planned trace), and their accuracy, i.e. whether the followed sequence was feasible or not.

#### 2.2.2 Control Condition

Our novel planning task mainly demands the execution of visuospatial planning function, but also requires visuospatial analysis and working memory to some extent (Wilson et al., 1996; Oosterman et al., 2013). In order to control confounding factors, a control task with all the cognitive and perceptual functions needed to solve the experimental task was designed, removing the component that elicits the planning function.

The control task had the same structure as the planning task. It consisted of the same 36 distinct mazes but each of them presented a traced path in a slightly darker color. These traced paths either follow the rules or not (see Figure 1B). Trials were pseudo-randomized. This task was also composed of three periods: control, execution, and response periods (see Figure 2B).

##### Control period (guided sequences)

Subjects were instructed to look at the mazes which had a traced path from the entrance visiting all four animals. Subjects had to evaluate the traced path and verify whether the sequence followed the rules or not. First, a fixation cross appeared for three seconds. The subject then had 10 seconds to evaluate the traced path. Same as the planning task, subjects could press a joystick button whenever they finished, and the reaction time was saved for further analyses.

##### Execution control period

A shifted fixation cross was presented for a three-seconds period just before the maze was presented again. The fixation cross preceded the location of the maze entrance to facilitate the gaze tracing. Subjects had 10 seconds to follow the traced path again, overlapping their gaze with the traced path. Once they reached the fourth animal they could finalize the trial by pressing a button and the reaction time here was recorded as well.

##### Response control period

During this period, a question mark appeared, and subjects were asked to answer whether the sequence was correct or not using joystick buttons and the accuracy response was saved. Finally, a feedback was presented, same as in the experimental condition.

For both planning and control conditions, subjects were orally instructed by the experimenter using visual aid before starting each task. Afterwards, a training session of six trials was held for each condition to ensure subjects got familiar with the experiment setup.

### 2.3. EEG Data Acquisition

Electroencephalography brain activity was recorded using a scalp EEG Biosemi® System (www.biosemi.com) consisting of sixtyfour scalp electrodes placed following the 10/20 system, and eight external electrodes. Four external electrodes measured electro-oculography (EOG) activity, two were used for electrocardiogram (EKG), and two for mastoids which were used for referencing later during signal pre-processing. All electrodes were placed according to standard anatomical references (Keil et al., 2014) and referenced to CMS and DRL active electrodes during acquisition. The data was sampled online with a rate of 2048 Hz.

### 2.4. Data Analyses

#### 2.4.1. Behavioral Data Analysis

Behavioral data were analyzed using custom scripts from MATLAB 8.0 (The Mathworks, Inc., Natick, Masachussets, United States), SPSS version 22 (IBM Corp. Released 2013. IBM SPSS Statistics for Windows, version 22.0. Armonk, NY: IBM Corp.), and GraphPad Prism version 6 for Windows (GraphPad Software, La Jolla California USA, www.graphpad.com). All behavioral analyses were conducted on the first two periods of each condition: planning, planning execution, control and control execution period.

The internal consistency of the task was evaluated by Cronbach’s Alpha coefficient. The following behavioral parameters were considered to analyze performance: accuracy (percentage rate of incorrect and correct responses) and reaction time (the average of time spent solving the mazes and evaluating marked paths, and all execution periods). D'Agostino & Pearson Omnibus Normality Distribution Test was conducted in order to choose the proper statistic test in each comparison (parametric or non-parametric). Furthermore, to evaluate homoscedasticity, the Levene Test was conducted. Depending on the data normality, Wilcoxon sign-ranked test or t-test matched-paired were performed to compare the difference between condition periods.

#### 2.4.2. Electrophysiological Data Analysis

##### 2.4.2.1. Signal Preprocessing

The EEG data pre-processing pipeline was carried out using EEGLAB toolbox codes (Delorme & Makeig, 2004), EYE_EEG extension (Dimigen, Sommer, Hohlfeld, Jacobs, & Kliegl, 2011), and the ADJUST plugin (Mognon, Jovicich, Bruzzone, & Buiatti, 2011).

Eye movement activity recorded from ET was synchronized with EEG recordings allowing to observe the occurrence of fixation, saccades, and blink events, improving the quality of the visual inspection. Co-registration was ensured with shared TTL trigger pulses that were sent from the presentation display PC to the ET computer during the whole experiment. The sampling rate was down-sampled to 1024 Hz and re-referenced to average of electrodes on mastoids. Then, a zero-phase finite impulse response (FIR) filter was used for high-pass filtering, with a high-pass cut-off frequency of 1 Hz and a low-pass cut-off frequency of 40 Hz. The EEG signal was segmented into 36 trials per condition, time-locked to the onset of planning and control periods as epochs of interest. Each trial consisted of 1 s before the start of the maze presentation (as a baseline) and 4 s after the planning or control period, respectively.

Subsequently, Logistic Infomax Independent Components Analysis (ICA) algorithm (Bell & Sejnowski, 1995) was used to identify and remove artefactual components from EEG data. Artefactual components associated with eye movements were rejected based on their covariance with simultaneously recorded eye movement data. This was done using saccade-to-fixation variance ratio criterion between 10 ms pre- and post-saccade (Plöchl, Ossandón, & König, 2012). Additionally, others artefactual components associated to EMG, electrode movement or non-brain-related components were identified by visual inspection. All rejected independent components were also visually validated by inspecting the topographies, spectra, and activations over time.

Finally, noisy channels identified by visual inspection and by automatic channel rejection using kurtosis criterion (5 z-score as threshold) were interpolated using spherical interpolation (typically Fp1 and Fp2).

##### 2.4.2.2. Time-Frequency Decomposition

EEG time-frequency analysis was carried out using short-time Fast-Fourier Transform (FFT) for frequencies ranging from 1-40 Hz using a window length of 250 ms and a time step of 5 ms. The time-frequency charts were then z-score normalized to the baseline (−1000 to −0.9 ms).

Topographic maps from the averaged theta frequency power of the whole epoch across the subjects in specific time points (750, 1750, 2750 and 3750 ms) during the 4000 ms of the trial were visualized. Electrodes Fz and Pz were selected for further analyses. Statistical comparisons of time-frequency charts from both conditions for Fz and Pz electrodes were made through a non-parametric cluster-based permutation test with p < 0.05. The statistic value chosen to perform the permutation test was the maximum statistic value of the cluster (Maris & Oostendveld, 2007).

Theta frequency band (4–8 Hz) was averaged along the whole trial (0 to 4 seconds) and then compared between conditions using a matched-pair t-test for electrodes Fz and Pz, respectively. To analyze the time profile of this band, power in the 4-8 Hz range was averaged across trials by subject. Time profiles of theta band activity for both conditions and electrodes were then compared using Wilcoxon Signed-Rank Test (match-paired, 88 ms steps of non-overlapping windows) and corrected by False Discovery Rate (FDR).

All time-frequency analyses were made using self-written scripts in MATLAB R2014a and R2018b and Statistics Toolbox 8.1 (The Mathoworks, Inc., Natick, Masachussets, United States).

##### 2.4.2.3. Source Reconstruction Analyses

Source localization analyses were performed using the open access Brainstorm toolbox (Tadel, Baillet, Mosher, Pantazis, & Leahy, 2011), which is documented and freely available for download online under the GNU general public license (http://neuroimage.usc.edu/brainstorm).

Sources were estimated over the preprocessed EEG signal (1-40 Hz range, filtered and cleaned) using Standardized Low-Resolution Brain Electromagnetic Tomography (sLORETA) (Pascual-Marqui, 2002). The parameters chosen to perform sLORETA were the minimum-norm imaging method, and the symmetric Boundary Element Method (symmetric BEM) using OpenMEEG toolbox (Gramfort et al., 2010). sLORETA algorithm was conducted on the default anatomical MNI template implemented in Brainstorm (“Colin27”) using the default electrode locations for each subject.

Consequently, the theta frequency band was selected because of the significance difference observed in the planning condition as compared to the control condition in time-frequency charts from Fz electrode. Also, theta band power increase in midline frontal electrodes was observed during visualization of topographic maps. Thus, source activity was band-pass filtered between 3.5 to 8.5 Hz. A z-score normalization was applied using −1000 to −10 ms pre-trial as baseline. Moreover, we selected a time span of interest since main FMθ activity started after 500 ms from time-frequency charts of Fz electrode and the temporal dynamics of topographic maps. Thus, source activity ranging from 3.5-8.5 Hz between 750 and 4000 ms was averaged. Subsequently, averaged space sources were compared between conditions using non-parametric permutation sign test using Monte-Carlo Sampling (1000 randomizations) (Tadel et al., 2011).

Cortical areas were labeled according to Destrieux Atlas available in the FreeSurfer Package in Brainstorm toolbox (Destrieux, Fischl, Dale & Halgren, 2010). Regions of interest (ROIs) were bilaterally selected based on significance differences between conditions in permutation tests as well as high theta frequency band increases in the planning condition alone. These regions included superior frontal gyri (SF), transverse frontopolar gyri and sulci (FP), anterior (ACC) and middle (mACC) part of the cingulate gyri and sulci (**Table 1**).

**Table 1.**
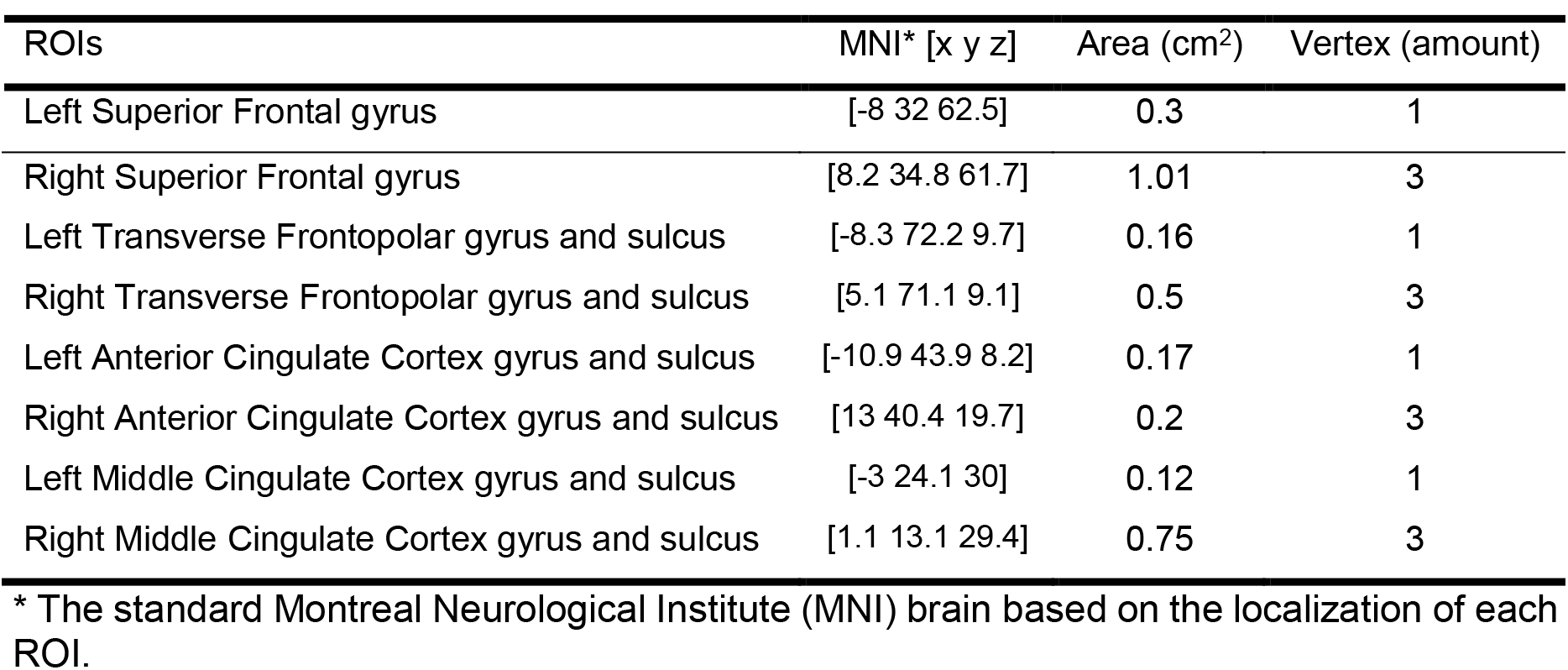
ROIs Brain Activity Sources.

| ROIs | MNI* [x y z] | Area (cm<sup>2</sup>) | Vertex (amount) |
| --- | --- | --- | --- |
| Left Superior Frontal gyrus | [-8 32 62.5] | 0.3 | 1 |
| Right Superior Frontal gyrus | [8.2 34.8 61.7] | 1.01 | 3 |
| Left Transverse Frontopolar gyrus and sulcus | [-8.3 72.2 9.7] | 0.16 | 1 |
| Right Transverse Frontopolar gyrus and sulcus | [5.1 71.1 9.1] | 0.5 | 3 |
| Left Anterior Cingulate Cortex gyrus and sulcus | [-10.9 43.9 8.2] | 0.17 | 1 |
| Right Anterior Cingulate Cortex gyrus and sulcus | [13 40.4 19.7] | 0.2 | 3 |
| Left Middle Cingulate Cortex gyrus and sulcus | [-3 24.1 30] | 0.12 | 1 |
| Right Middle Cingulate Cortex gyrus and sulcus | [1.1 13.1 29.4] | 0.75 | 3 |

\* The standard Montreal Neurological Institute (MNI) brain based on the localization of each ROI.

A second source analysis was conducted over the preprocessed EEG signal (1-40 Hz range, filtered and cleaned) following the same procedure explained above except for the application of a band-pass filter between 3.5 to 8.5 Hz. Thereafter, Principal Component Analysis (PCA) was conducted for each ROI’s activity and the first mode of the PCA decomposition for each ROI was selected. A spectral estimation using Morlet Wavelet Transform was performed on these selected components. In a linear frequency definition from 1-25 Hz in 1 by 1 frequency step, 1 Hz as central frequency and 3 s time resolution were selected. To avoid edge effects, each end of the signal was mirrored using a length of 512 samples. The frequency power was normalized to z-score using −1000 to −10 ms as baseline. Time-frequency charts were compared between periods using non-parametric cluster-based permutation tests (Maris & Oostendveld, 2007).

To compare the time profile of theta band across conditions, activity from selected ROIs was band-pass filtered between 4-8 Hz and Hilbert Transform was applied to obtain the instantaneous amplitude (Le Van Quyen et al., 2001). Here as well, to avoid edge effects, each end of the signal was mirrored using a length of 512 samples. Then, the signal was z-scored (using −1000 to −10 ms as baseline), down-sampled to 125 Hz, and averaged across trials by subject. Each ROI source activity was compared between conditions using Wilcoxon Signed-Rank Test (matched-pairs, 500 or 750 ms steps of non-overlapping windows) and corrected by FDR. Morlet Wavelets and Hilbert Transform were applied, and envelope calculations were performed using Complex Wavelet Toolbox and Signal Processing Toolbox from Matlab software (MathWorks).

## 3. Results

### 3.1. Behavioral Results

All behavioral features analyzed showed a normal distribution according to D'Agostino & Pearson Omnibus Normality Test except values of accuracy for control condition. Supplementary **Table S1** shows a summary of normality distribution test results.

#### 3.1.1. Reliability

The internal consistency of each task period (planning, control, execution planning, execution control) was excellent according to the categories of reliability proposed by George and Mallery (2003). The Cronbach’s alpha coefficient ranged between 0.95 and 0.97 (Supplementary **Table S2**) for reaction times as input. These results suggest that each task period has a consistent set of trials indicating a high task reliability.

#### 3.1.2. Performance

The **Table 2** shows a summary of the most relevant descriptive statistics.

**Table 2.** Behavioral descriptive data for each parameter.

| Parameters | Descriptive Statistics* |  |  |  |
| --- | --- | --- | --- | --- |
|  | Mean | Median | SD | SEM |
| Planning RT | 8.85 | 9.15 | 1.15 | 0.22 |
| Execution Planning RT | 6.33 | 6.29 | 1.11 | 0.21 |
| Accuracy Planning | 85.70 | 88.89 | 7.69 | 1.48 |
| Control RT | 6.81 | 6.57 | 1.50 | 0.29 |
| Execution Control RT | 5.71 | 5.55 | 0.99 | 0.19 |
| Accuracy Control | 95.68 | 97.22 | 4.77 | 0.92 |

SD = Standard deviation; SEM = Standard error of the mean; RT = Reaction Time
\*Values of RT are presented in seconds and values of accuracy are percentage of correct responses.

Variability analyses by Levene Test showed homogeneity in variance (Supplementary **Table S3**).

The mean reaction time of the planning period in comparison to control period was significantly greater (**Table 2** **and** **3**, **Figure 3A**). The same was observed when the reaction time of the execution planning period was compared to the execution control period (**Table 2** **and** **3**, **Figure 3B**). This reflected that the planning condition was cognitively more demanding than the control condition.

**Table 3.** Behavioral Performance Statistical Comparison.

| Parameters* |  | Statistical hypothesis |  |
| --- | --- | --- | --- |
|  |  | t-test / Wilcoxon | p-value |
| RT Planning | RT Control | t (26) = 6.23 | <0.0001**** |
| RT Execution Planning | RT Execution Control | t (26) = 3.41 | 0.0021** |
| RT Planning | RT Execution Planning | t (26) = 7.65 | <0.0001**** |
| RT Control | RT Execution Control | t (26) = 4.02 | 0.0004*** |
| Accuracy Planning | Accuracy Control | Z = 375 | <0.0001**** |

\*Values of RT used were in seconds and values of accuracy were percentage of correct responses.

**Figure 3.**
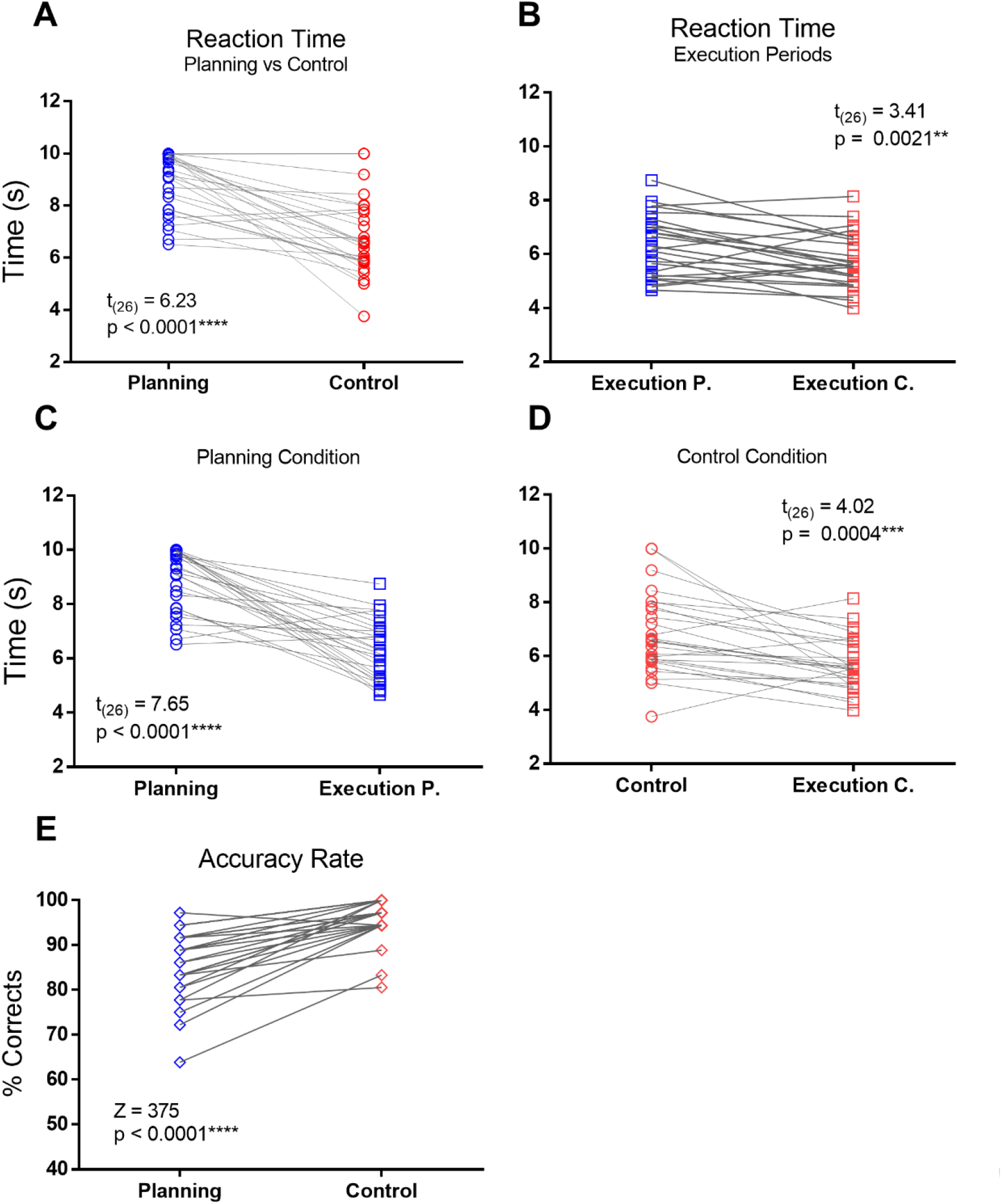
Behavioral Performance for Planning condition (in blue) and Control condition (in red). Comparison between (A) average reaction time in planning period (blue circles) and control period (red circles); (B) average reaction time in execution planning period (blue squares) and execution control period (red squares); (C) average reaction times in planning period (blue circles) and execution planning period (blue squares); (D) average reaction times in control period (red circles) and execution control period (red squares); (E) accuracy rate in planning condition (blue diamonds) and control conditions (red diamonds).

Furthermore, the planning period was also more time consuming than the execution planning period (**Table 2** **and** **3**, **Figure 3C**). Similarly, the reaction time during the control period was significantly greater than the reaction time of the execution control period (**Table 2** **and** **3**, **Figure 3D**).

In terms of response accuracy, subjects were less accurate during the planning condition as compared to the control condition (**Table 2** **and** **3**, **Figure 3E**) which may also reflect that the planning condition is more complex leading the subjects to perform less accurately.

Collectively, once the planning component was successfully extracted from the control condition, these behavioral results indicate that the planning condition (both the planning and the execution planning period) is more cognitive demanding. This was reflected by high response time and less accuracy during the planning condition. The task conditions, specifically their neural correlates, can also be compared with each.

### 3.2. Electrophysiological Results

Topographic maps corresponding to the average theta band power during 4 seconds of planning period showed a local increase in the frontal midline electrodes (Fpz, AFz, Fz, FCz) as compared to control period where there was not an apparent increase in theta band power. The planning effect was computed as the power subtraction between planning and control periods and showed an increase in theta frequency band that stays consistent in frontal midline electrodes from the planning period (**Figure 4**).

**Figure 4.**
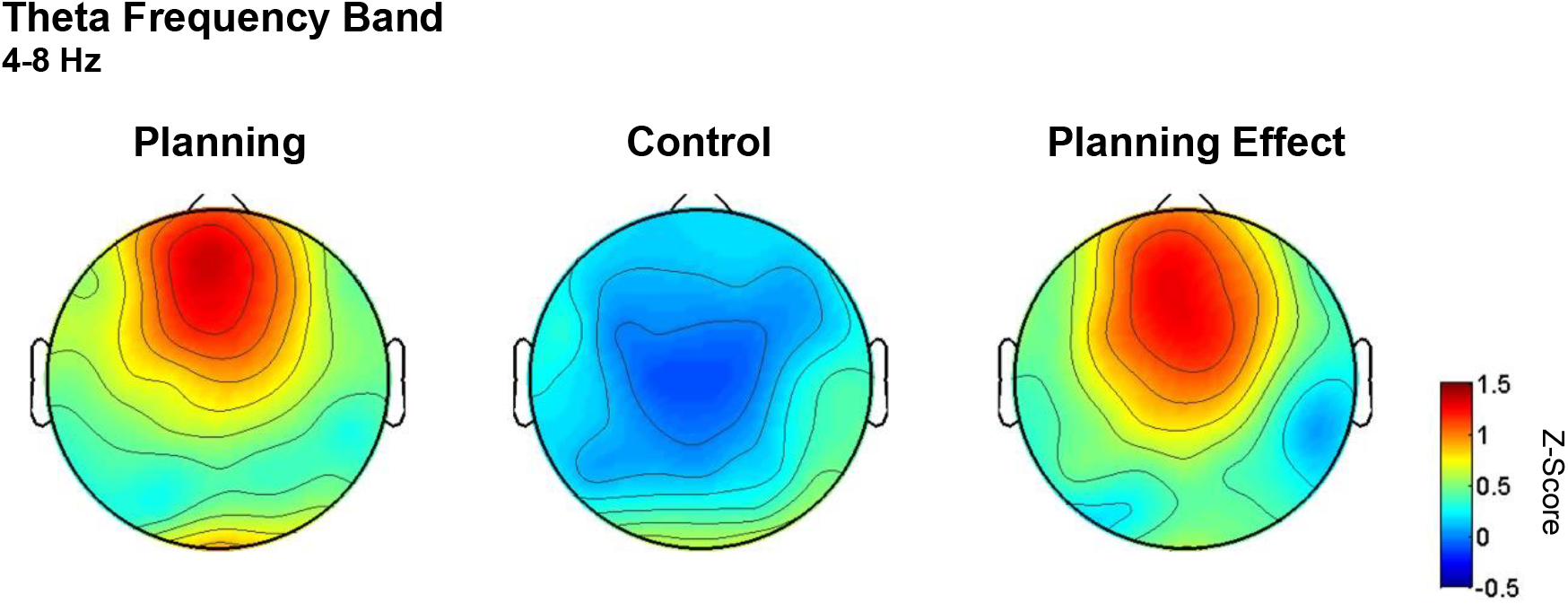
Topographic maps of average theta band power during the first 4 seconds of planning. Scalp representation of average theta frequency band power across all subjects, normalized to z-score during planning period (left), control period (middle) and for the planning effect (right). Left: During planning period, subjects showed an increase in theta frequency band power located mainly in frontal midline electrodes (Fpz, AFz, Fz, FCz). Middle: Control period showed no evident increase in theta frequency band, except for posterior (occipital) electrodes. Right: Planning Effect (power subtraction between periods: Planning – Control) illustrates the increase in theta frequency band in frontal midline electrodes.

Time frequency analysis of electrodes Fz and Pz revealed a marked increase in low frequency bands, most importantly a progressive increase in the theta band (4-8 Hz) that starts at 1133.8 ms of planning period onset (**Figures 5**–**6**, **Table 4**). This effect was completely absent in the control period. This difference was assessed using a cluster-based permutation test (**Figure 5**) confirming a significant difference in the theta band power between planning and control periods for both Fz and Pz electrodes. Furthermore, a significant power decrease was found in both alpha and beta ranges during the planning condition when compared to the control condition (**Figure 5**, **Table 4**).

**Table 4.** Cluster-based Time-Frequency Comparison.

| Fz Electrode |  |  |  |  |  |
| --- | --- | --- | --- | --- | --- |
| Positive Cluster # | p-value | Cluster-stat | SD | Time Window<br>(ms) | Frequency<br>Range (Hz) |
| 1 | 0.0020 | 14209.9 | 0.0014 | 1133.8 – 3877.9 | 1 – 8 |
| Negative Cluster # |  |  |  |  |  |
| 1 | 0.0030 | -9773,4 | 0.0017 | 1319.3 – 2706.1 | 18 – 32 |
| 2 | 0.0040 | -9053,7 | 0.0020 | 2769.5 - 3877.9 | 18 – 38 |
| 3 | 0.0090 | -5937,9 | 0.0030 | 1.0 – 669.9 | 10 – 40 |
| Pz Electrode |  |  |  |  |  |
| Positive Cluster # | p-value | Cluster-stat | SD | Time Window<br>(ms) | Frequency<br>Range (Hz) |
| 1 | 0.0120 | 4737.5 | 0.0034 | 1133.8 – 2945.3 | 1 – 7 |
| Negative Cluster # |  |  |  |  |  |
| 1 | 0.0040 | -9022.7 | 0.0020 | -384.8 – 635.7 | 4 – 40 |
| 2 | 0.0130 | -4769.0 | 0.0036 | 2813.5 – 3877.9 | 9 – 28 |

SD = Standard deviation ; ms = milliseconds ; Hz = Hertz.

**Figure 5.**
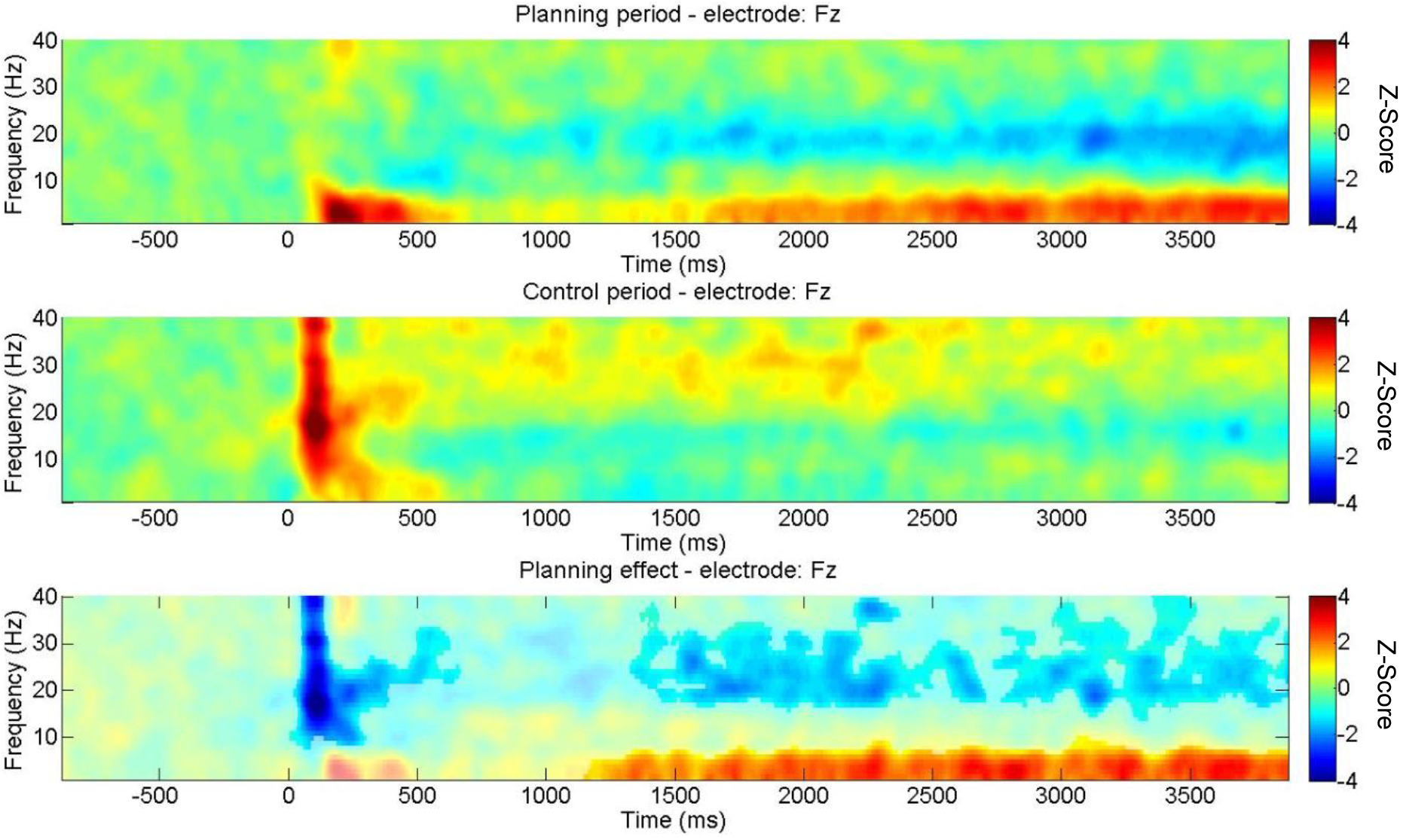
Time-Frequency charts for Fz electrode. Top: Time-frequency plot for the planning period showing a marked increase in theta band power with time. Middle: Time-frequency plot for control period. Bottom: Planning effect computed as the subtraction between planning and control period. Non-significant pixels are shown lighter in the plot.

**Figure 6.**
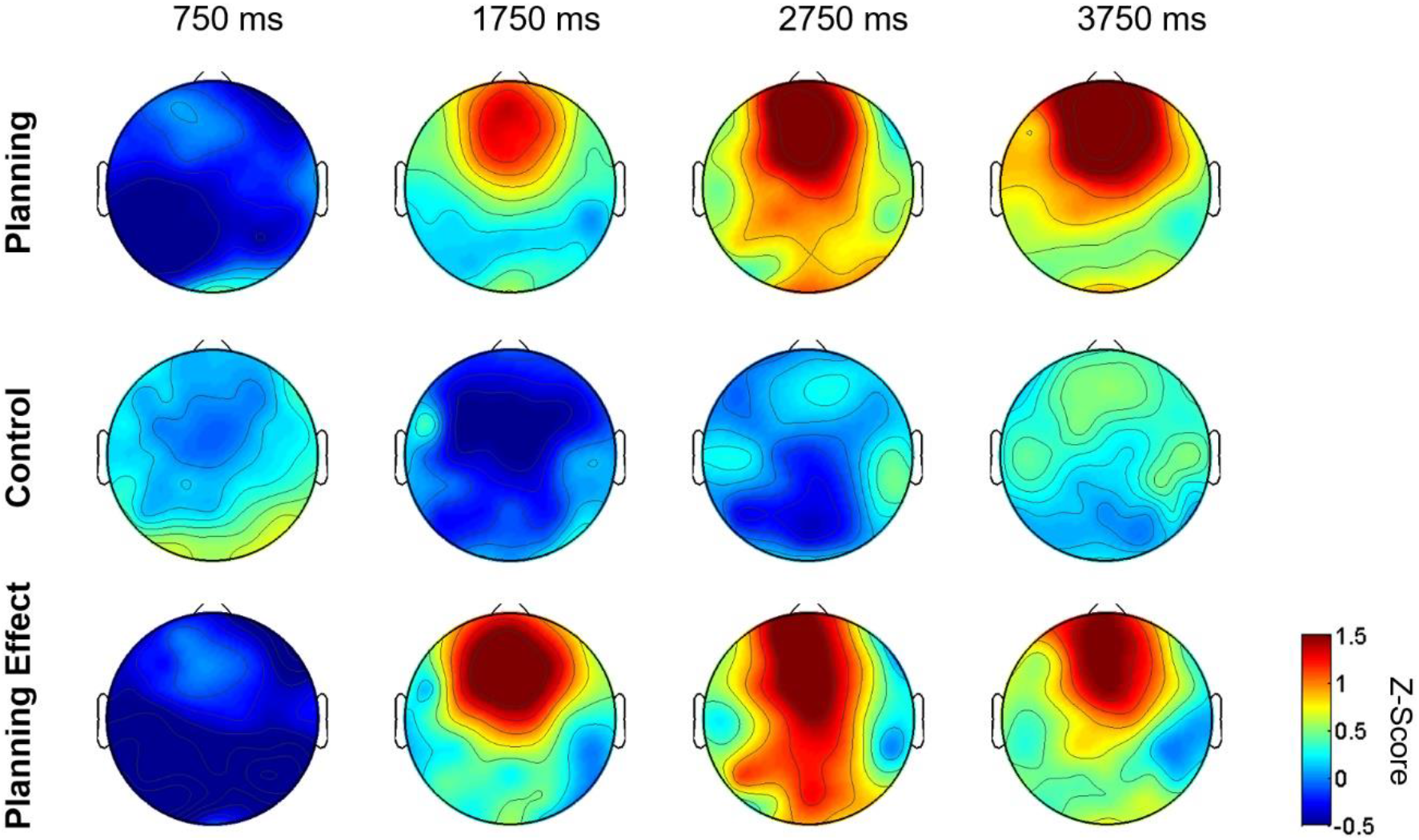
Theta temporal dynamic of topographic maps. Different time points are depicted in topographic scalp maps. We observed a progressive increase in frontal midline theta frequency band across time while subjects were planning paths and not when they were evaluating them. Each topographic map shows theta frequency power normalized to z-score averaged over all 36 trials for all 27 subjects at a specific time point. Color bar represents z-score between the values of −0.5 (blue) to 1.5 (red).

In order to better characterize the increase in theta frequency band, we then averaged the power between 4-8 Hz for both periods, obtaining the average band power of theta band over time. We found that increase in theta activity was significantly greater for planning period for Fz but not for Pz electrodes (**Figure 7**, Supplementary **Table S4**).

**Figure 7.**
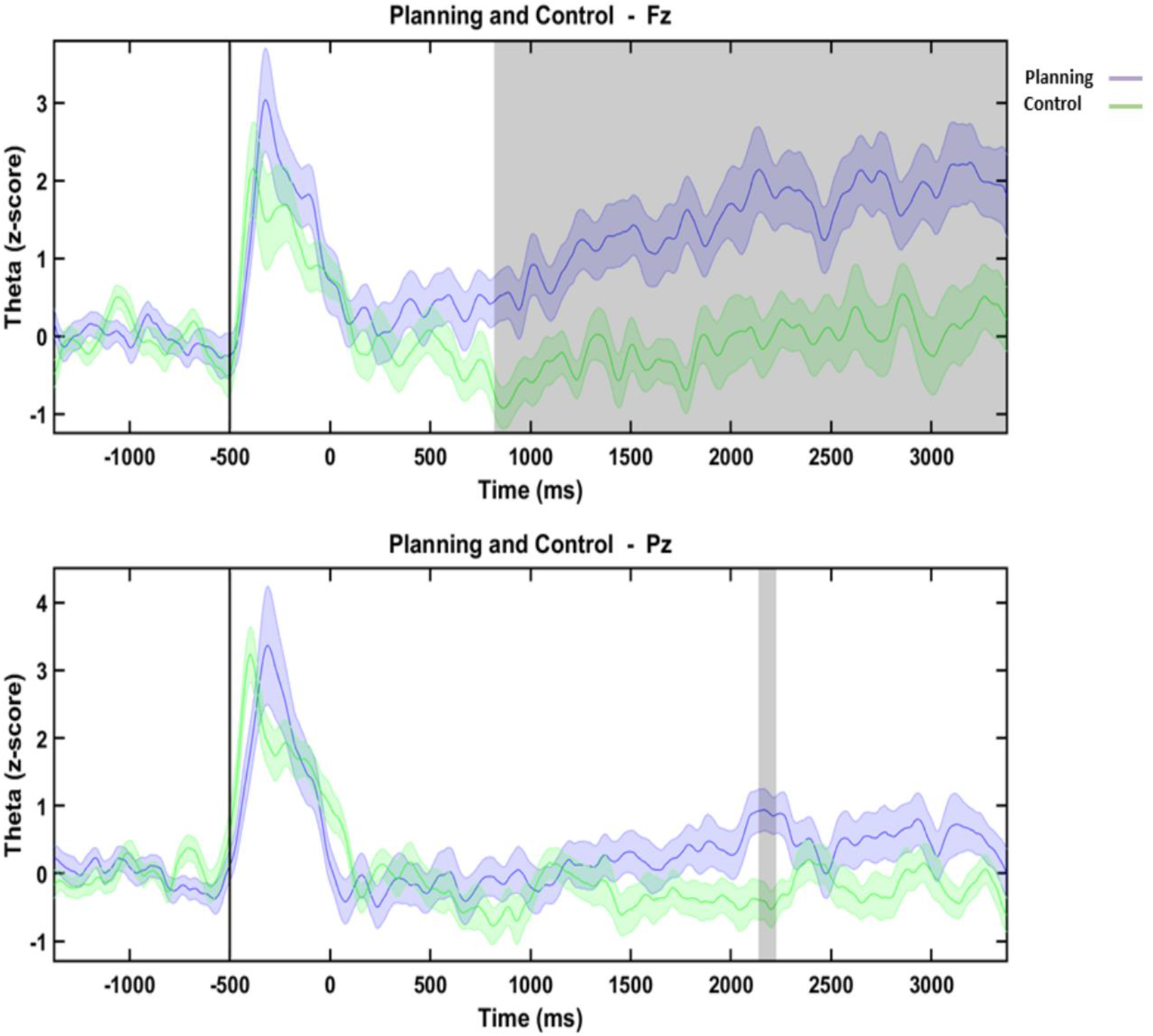
Time Theta Frequency Slices. Fz (top) and Pz (bottom) electrodes and their averaged theta band power over time for the planning period (in purple) and the control period (in green). Both electrodes and periods exhibited a theta increase at the beginning of the trial (before 500 ms) and then, only Fz electrode for planning period showed a sustained increase in theta band activity over time (Wilcoxon test, FDR corrected, see **Table 7**). Gray shaded areas show statistically significant differences according to a non-overlapping moving window with steps of 88 ms of Wilcoxon signed-rank test comparison. Shaded purple and green color around theta power mean of each condition corresponds to standard error of the mean.

To analyze global theta activity during the whole epoch, the averaged theta frequency band during the first 4 seconds of planning for Fz from planning period (mean = 1.04, SD = 1.26, SEM = 0.24) was compared to control period (mean = 0.04, SD = 0.86, SEM = 0.17, **Table 7**). Global theta activity from Fz electrode during planning showed a significant increase in comparison to global theta activity during the control period (paired-samples t-test, t (26) = 3.86, p = 0.0007***, **Figure 8A**, **Table 5**). However, no significant differences between conditions were found in the case of Pz (planning period, mean = 0.34, SD = 1.00, SEM = 0.19; control period, mean = 0.04, SD = 0.53, SEM = 0.10; paired-samples t-test, t (26) = 1.50, p = 0.15, **Figure 8B**, **Table 5**).

**Table 5.** Descriptive Statistics of Averaged Theta Activity.

| Electrode and Period | Descriptive Statistics* |  |  | Normality Distribution<br>D'Agostino & Pearson omnibus |  |
| --- | --- | --- | --- | --- | --- |
|  | mean | SD | SEM | K2 | P value |
| Fz Planning | 1.04 | 1.26 | 0.24 | 0.67 | 0.72 |
| Fz Control | 0.04 | 0.86 | 0.17 | 5.94 | 0.05 |
| Pz Planning | 0.34 | 1.00 | 0.19 | 0.68 | 0.71 |
| Pz Control | 0.04 | 0.53 | 0.10 | 1.67 | 0.43 |

\*Values are z-score units; SD = Standard deviation; SEM = Standard error of the mean.

**Table 6.** Cluster-based Time-Frequency Comparison of Sources.

| Cluster # | p-value | Cluster-stat | SD | Time Window<br>(s) | Frequency<br>Range (Hz) |
| --- | --- | --- | --- | --- | --- |
| **Frontal Superior L** |  |  |  |  |  |
| Positive Cluster 1 | 0.0020 | 9229.4 | 0.0014 | 0.0 – 1.2 | 2 – 6 |
| Positive Cluster 2 | 0.0070 | 7487.7 | 0.0026 | 2.0 – 4.0 | 2 – 6 |
| **Frontal Superior R** |  |  |  |  |  |
| Positive Cluster 1 | 0.0040 | 6960.4 | 0.0020 | 0.0 - 1.0 | 2 – 6 |
| **Frontopolar L** |  |  |  |  |  |
| Positive Cluster 1 | 0.0250 | 5409.6 | 0.0049 | 1.5 – 2.4 | 3 – 6 |
| Positive Cluster 2 | 0.0260 | 5306.2 | 0.0050 | 0.1 – 1.0 | 2 – 6 |
| **Frontopolar R** |  |  |  |  |  |
| Positive Cluster 1 | 0.0270 | 4469.0 | 0.0051 | 1.6 – 2.4 | 4 – 6 |
| **mACC R** |  |  |  |  |  |
| Negative Cluster 1 | 0.0420 | -4092.2 | 0.0063 | 1.7 – 2.0 | 9 – 25 |

SD = Standard deviation ; ms = milliseconds ; Hz = Hertz.

**Table 7.** Time Source Theta Activity: Statistics Analyses Over Time.

| Superior Frontal L |  |  |  |  | Superior Frontal R |  |  |  |  |
| --- | --- | --- | --- | --- | --- | --- | --- | --- | --- |
| Time Window (ms) |  | Wilcoxon Signed-Rank Test |  |  | Time Window (ms) |  | Wilcoxon Signed-Rank Test |  |  |
| Start | End | p-value | FDR Threshold | z-value | Start | End | p-value | FDR Threshold | z-value |
| 0 | 768 | 0.0102 | 1 | 2.57 | 0 | 512 | 0.3613 | 0 | 0.91 |
| 768 | 1536 | 0.0881 | 0 | 1.71 | 512 | 1024 | 0.0163 | 1 | 2.40 |
| 1536 | 2304 | 0.0088 | 1 | 2.62 | 1024 | 1536 | 0.1635 | 0 | 1.39 |
| 2304 | 3072 | 0.0837 | 0 | 1.73 | 1536 | 2048 | 0.0109 | 1 | 2.55 |
| 3072 | 3840 | 0.0271 | 1 | 2.21 | 2048 | 2560 | 0.1864 | 0 | 1.32 |
|  |  |  |  |  | 2560 | 3072 | 0.0133 | 1 | 2.47 |
|  |  |  |  |  | 3072 | 3584 | 0.0366 | 0 | 2.09 |

| Frontopolar L |  |  |  |  | Frontopolar R |  |  |  |  |
| --- | --- | --- | --- | --- | --- | --- | --- | --- | --- |
| 0 | 752 | 0.0271 | 0 | 2.21 | 0 | 752 | 0.0198 | 1 | 2.33 |
| 752 | 1504 | 0.0546 | 0 | 1.92 | 752 | 1504 | 0.2205 | 0 | 1.23 |
| 1504 | 2256 | 0.0071 | 1 | 2.69 | 1504 | 2256 | 0.0077 | 1 | 2.67 |
| 2256 | 3008 | 0.2488 | 0 | 1.15 | 2256 | 3008 | 0.1709 | 0 | 1.37 |
| 3008 | 3760 | 0.4140 | 0 | 0.82 | 3008 | 3760 | 0.6480 | 0 | 0.46 |

| ACC L |  |  |  |  | mACC L |  |  |  |  |
| --- | --- | --- | --- | --- | --- | --- | --- | --- | --- |
| 0 | 496 | 0.0461 | 0 | 1.99 | 0 | 504 | 0.2391 | 0 | 1.18 |
| 496 | 992 | 0.4140 | 0 | -0.82 | 504 | 1008 | 0.0066 | 1 | 2.71 |
| 992 | 1488 | 0.0116 | 1 | 2.52 | 1008 | 1512 | 0.4860 | 0 | 0.70 |
| 1488 | 1984 | 0.0109 | 1 | 2.55 | 1512 | 2016 | 0.0926 | 0 | 1.68 |
| 1984 | 2480 | 0.1023 | 0 | 1.63 | 2016 | 2520 | 0.2297 | 0 | 1.20 |
| 2480 | 2976 | 0.5642 | 0 | 0.58 | 2520 | 3024 | 0.0837 | 0 | 1.73 |
| 2976 | 3472 | 0.7186 | 0 | 0.36 | 3024 | 3528 | 0.1363 | 0 | 1.49 |
| 3472 | 3968 | 0.0152 | 1 | 2.43 |  |  |  |  |  |

FDR = False Discovery Rate; 0 = False; 1 = True.

**Figure 8.**
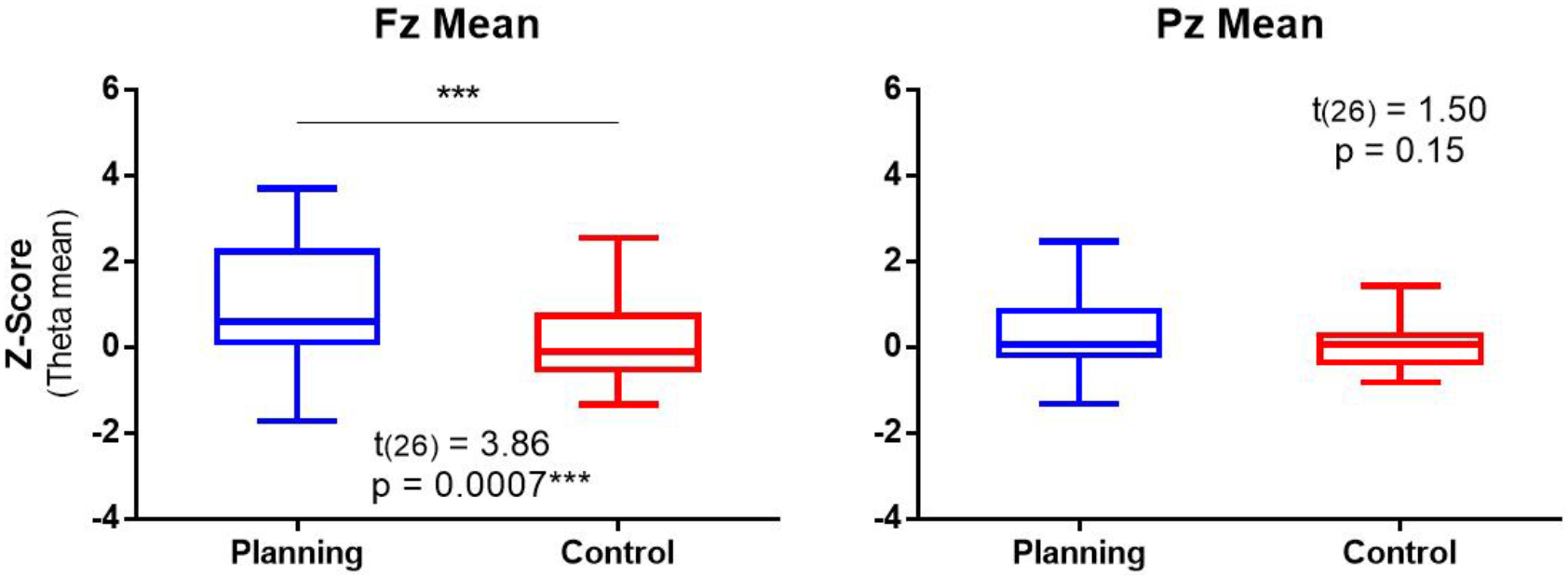
Global Theta Activity. Fz average theta band power (left) for all 27 subjects during the first 4 seconds of planning (blue) in comparison to the control period (red) exhibited a significant greater power (paired-samples t-test, t (26) = 3.86, p = 0.0007***). In contrast, Pz (right) did not present significant differences between periods (t-test paired samples, t (26) = 1.50, p = 0.15). The hinges of the box extend from the 25th to 75th percentiles. The line in the middle represents the median. The whiskers show the smallest and the largest value. Descriptive statistics in **Table 9**.

### 3.3. Source Reconstruction

Sources were estimated using sLORETA (Pascual-Marqui, 2002) across the epoch for the 3.5-8.5 Hz frequency range, then z-score normalized (using 1 second pre-trial as baseline) and averaged between 0.750 and 4 seconds. This span of time was selected based on the significant differences showed in the time profile of theta increase for Fz electrode. In order to localize the sources, we visualized the whole brain model template and cortical activations for both conditions.

We found a specific activation in prefrontal areas for planning, and right-occipital and right-temporal activations for the control period. However, significant differences were found in PFC (bilateral SF and ACC, **Figure 9)**.

**Figure 9.**
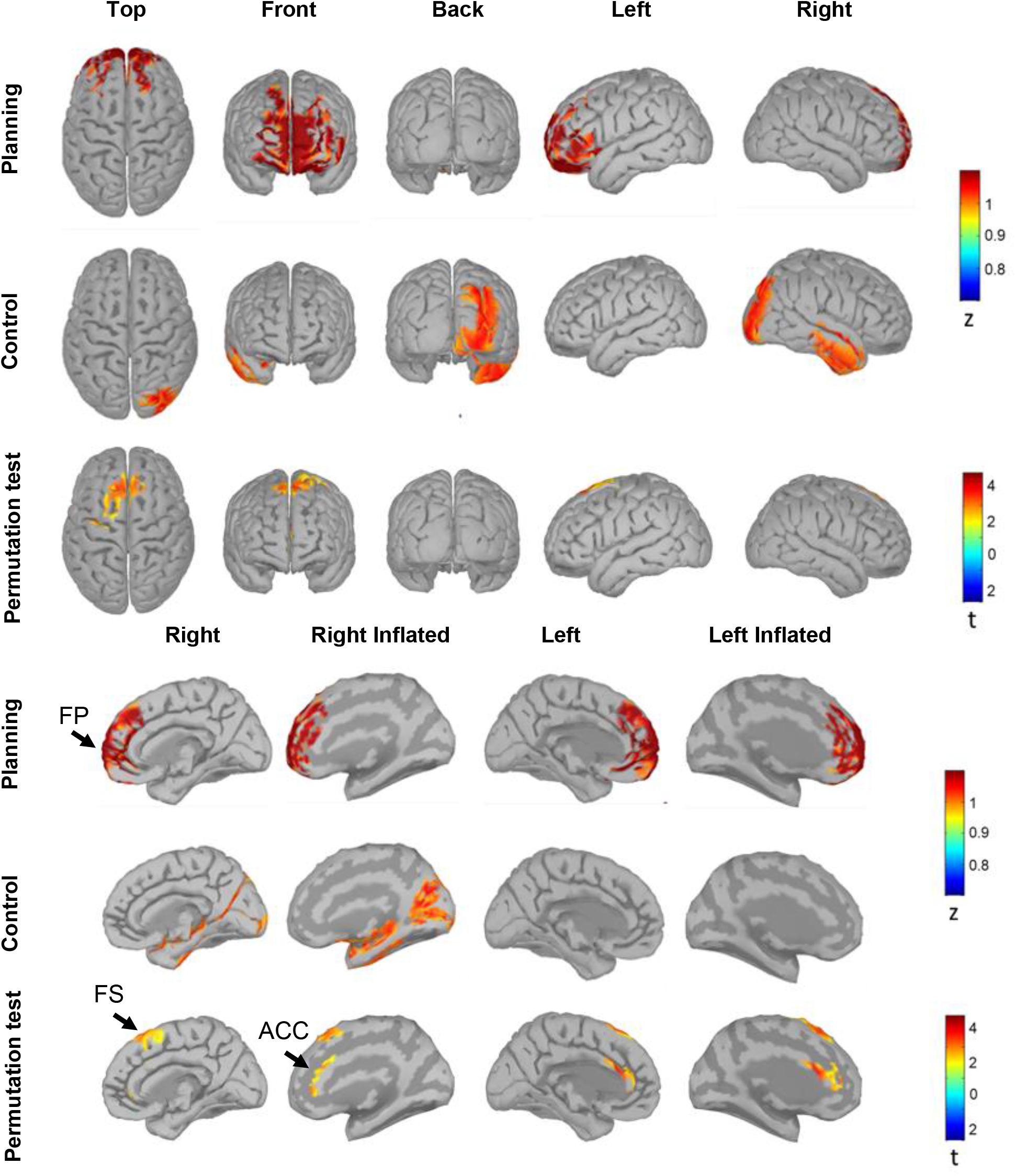
Source Reconstruction. sLORETA was applied to the task epoch for the preprocessed EEG signal (3.5-8.5 Hz bandpass filtered) for both conditions, normalized to z-score and averaged between 0.75 and 4 seconds. Sources were projected to a template brain model. On the top, a whole brain model is shown. At the bottom, a sagittal plane of the brain is shown. Visualization of sources showed increases in theta activity in lateralized right occipital and temporal regions for the control condition. The increase in theta activity in prefrontal region and anterior cingulate cortex is noticeable in the planning period. The plot shows absolute values of z-score between 0.77 and 1.08. Black arrows show FP, FS and ACC labels. FP: Frontopolar; FS: Frontal Superior; ACC: Anterior Cingulate Cortex.

Furthermore, to analyze the time frequency domain we performed spectral estimation of ROI time series. ROI time series were calculated using the first mode of the PCA decomposition of all the signals from a ROI. Time-frequency charts were then obtained by using Morlet Wavelet Transform. Subsequently, time-frequency charts were compared between periods using non-parametric cluster-based permutation tests (**Figure 10**, **Table 6**). Bilateral FS and FP sources presented significant positive clusters in 2-6 Hz range which includes the theta activity.

**Figure 10.**
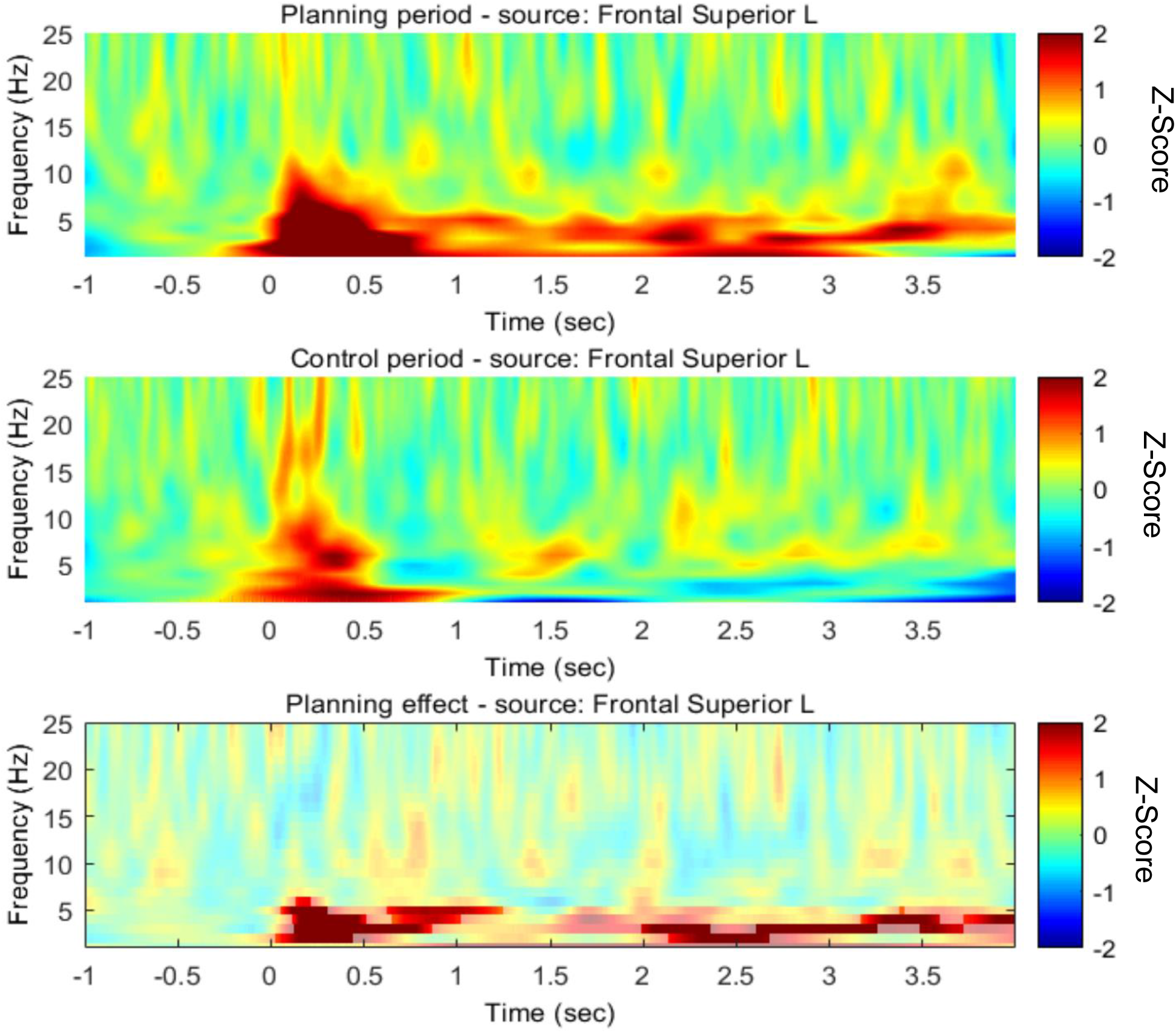
Time-Frequency charts for Left Superior Frontal Source. These Time-Frequency charts show averaged spectrograms across subjects over time for Left Superior Frontal source during the planning period (top) and the control period (middle). Cluster-based permutation test comparison between those conditions is showed at the bottom. Both periods showed increases in theta band power at the beginning of the trial which was greater for the planning period. 1 second after the planning period onset, a later theta activity was observed. Cluster-based permutation test found 2 main clusters, once during early stage and the next during the later stage of planning. Non-significant pixels are shown lighter in the chart. L = left.

Bilateral FS and right FP sources showed an early first cluster that appeared at the beginning of planning and finalized at about 1 second. A later positive cluster emerged and sustained between 1.5 and 2.4 s in the left FP, between 1.6 and 2.4 s in the right FP, and between 2 and 4 s in the left SF. Additionally, a negative cluster was found in the right mACC that was sustained between 1.7 and 2 seconds of planning in a broadband range of 9–25 Hz which includes alpha and beta frequency band (**Table 6**).

Finally, to evaluate theta changes over time, we performed a Hilbert Transform for each ROI time series after which we compared the amplitude of theta frequency between conditions. We found that bilateral SF presented significantly higher theta frequency band power after first 1.5 seconds of planning period onset. This lasted until 2.75 seconds for the left SF, and 2 seconds for the right SF. This also occurred after 3 until 4 seconds in the left SF, and after 2.75 until 3.25 in the right SF. The left SF also showed significant theta increases in the first 750 ms of planning period onset. Bilateral FP showed theta increases between 1.5 and 2.250 seconds of planning period onset. The right FP presented increases in theta power during the first 750 ms of planning period onset. Additionally, left ACC presented significant theta increases between 1 and 2 seconds, and between 3.5 and 4 seconds after planning period onset. Also, the left mACC showed theta increases between 0.5 and 1 second after planning period onset (**Figure 11**, **Table 7**).

**Figure 11.**
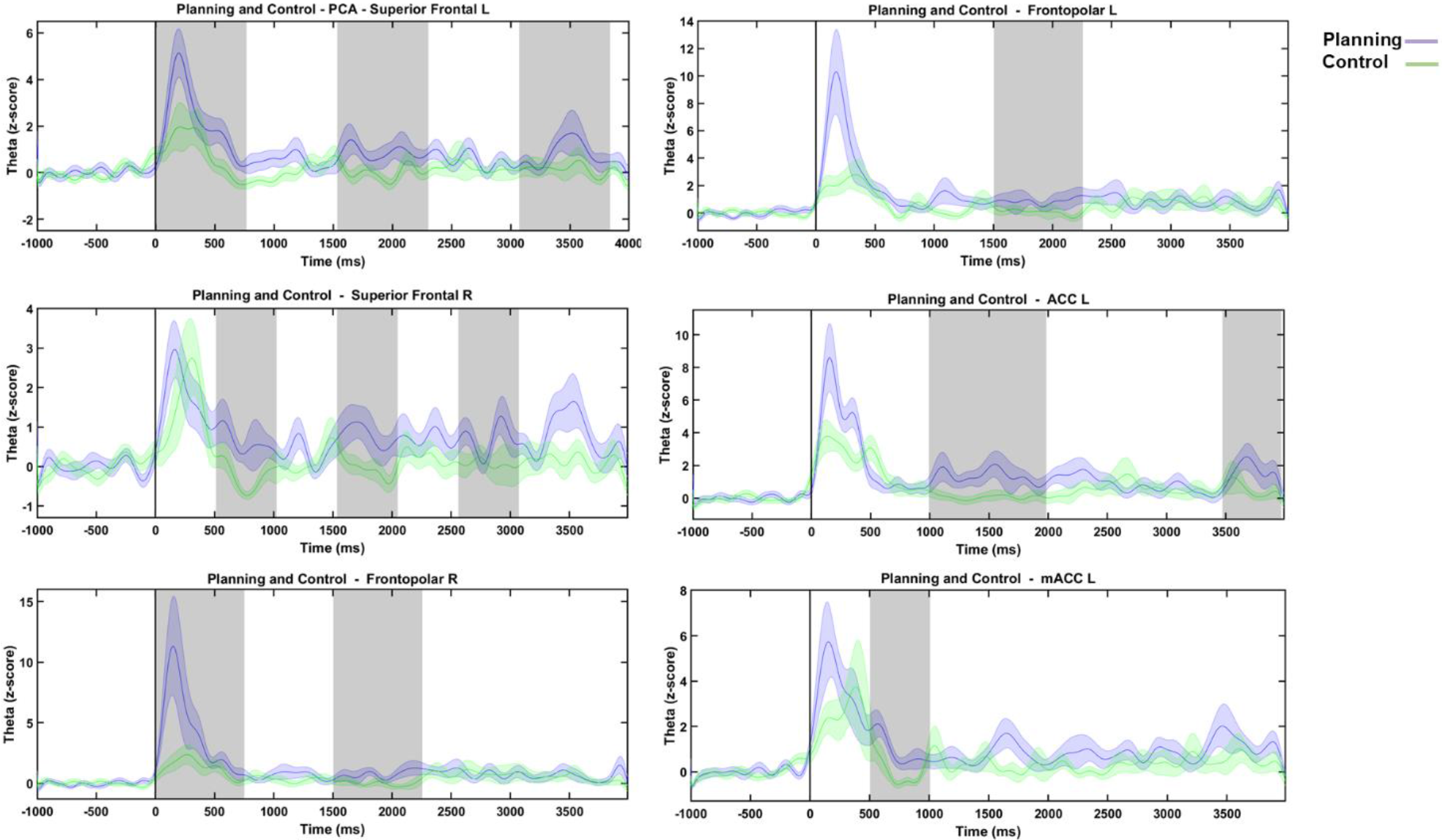
Source Theta Activity over Time. Hilbert Transform was applied to first mode of the PCA decomposition for each ROI (3.5-8.5 Hz bandpass filtered) and for both conditions (planning in purple, control in green), normalized to z-score, showing the instantaneous amplitude of source theta activity over time. Gray shaded areas show statistically significant differences according to a non-overlapping moving window with steps of 500 or 750 ms of Wilcoxon signed-rank test comparison. Shaded purple and green colors around theta power mean of each condition correspond to standard error of the mean. All ROI sources showed increases in theta activity at the beginning (at about 750 ms). Nevertheless, for the planning period, only the left Superior Frontal and the right Frontopolar areas showed higher activity during this early period. A later increase in theta amplitude for planning period was observed for all sources as seen in the shaded gray areas (Wilcoxon test, FDR corrected, see Table 11). L = Left; R = Right; ACC = Anterior Cingulate Cortex; mACC = middle Anterior Cingulate Cortex.

## 4. Discussion

In this study, we recorded EEG activity during an ecological planning task to evaluate whether cognitive planning, as a higher order cognitive control function, induces FMθ activity. To address this problem, we designed a novel planning task with adequate psychometric properties in terms of reliability and variability.

There are studies that have assessed the reliability of the outcomes generated from planning tasks (Wilson et al., 1996; Porteus, 1959), however, there have not been any reports on psychometric properties needed for the adaptation of a planning tasks for neuroimaging assessment (Kirsch et al., 2006; Tremblay et al.,1994) at least within the scope of our literature review. In this study excellent reliability was found for the behavioral task outcomes in the two conditions: planning and control (**Table S2**). Additionally, the behavioral parameters of the task showed a normal distribution (**Table S1**) and variance homogeneity (**Table S3**).

Designing an ecological behavioral task primarily requires predictive validity, i.e. that the task must be able to identify impaired planning function in patients with psychiatric or cognitive disorders who actually exhibit impaired planning performance in their daily life (Oosterman, Wijers, & Kessels, 2013). For this reason, our novel paradigm was based on the Zoo Map Task, which has been shown to have optimal predictive validity in previous studies (Oosterman, Wijers, & Kessels, 2013). Secondly, an ecological task design requires the paradigm to demand subjects to exert cognitive planning that an actual daily life situation would demand (Miotto & Morris, 1998; Burgess, Simons, Coates & Channon, 2005; Morris & Ward, 2005). In order to achieve this, our task considers a period in the beginning where subjects have to plan a path monitoring whether it follows the given set of rules, and then store it in working memory. Subsequently, there is a second period in which subjects must carry out their former plan while monitoring the behavioral execution of the path, making sure the path follows these rules. While executing the plan, subjects must have enough cognitive flexibility to adequately correct the trajectory if it is traced incorrectly. Thus, the task also demands attentional control as it may occurs in real-life situations, where high order cognitive functions are exerted in a concerted manner. Therefore, designing an adequate control task that effectively isolates the planning component was essential, and this is reflected in the results obtained in both behavioral and electrophysiological measures. Since, in control condition, subjects were asked to perform a goal which did not consist of planning but consisted of virtually the same stimuli, the planning component was successfully removed in the condition. Due to this removal, we observed FMθ activity for the planning period, which was not observed in the control period.

The behavioral results for the planning condition were in line with our predictions. Since the planning task implies a high cognitive function (Lezak, 1995; Zwosta, Ruge & Wolfensteller, 2015), we predicted higher time responses and less accurate performances during the planning condition (complex task: plan a path in a complex map) than during the control condition (simple task: they only have to follow a marked path and decide whether it followed the rules). Moreover, in both periods of the planning condition (the planning period and the n execution planning period) reaction times were always greater than the control condition (the control period and the execution control period) reflecting how difficult and cognitively demanding the planning condition is (Owen et al., 1996; Voytek et al, 2015; Ossandón et al, 2012). Interestingly, the execution of the planned path (during the execution planning period) involved a greater cognitive effort during the planning task, as suggested by higher reaction times during the planning execution period compared with the control condition periods (the control period and the control execution period). This can be explained by the requirement of high cognitive functions such as working memory and attentional control to perform the execution of the plan. All these observations are in line with the theoretical assumption of cognitive planning (Hayes-Roth & Hayes-Roth, 1979; Wilensky, 1983; Grafman & Hendler, 1991; Zwosta, Ruge & Wolfensteller, 2015).

Previous studies have reported that PFC has a critical role during cognitive planning (Kirsch et al., 2006; Newman, Carpenter, Varma, & Just, 2003; Owen et al., 1996; Nitschke et al., 2017) and the present results show that cognitive planning induces a FMθ activity (**Figures 4**-**8**) originating in PFC, specifically SF and ACC (**Figure 9**). These results are in line with previous studies on higher order cognitive functions (Cavanagh & Frank, 2014). Extensive evidence supports the role of FMθ activity as a common top-down mechanism for realizing the need of cognitive control (Cavanagh & Frank, 2014). FMθ activity, as a marker of cognitive control, is thought to be exerted by recruiting and aiding communication between brain regions during tasks that requires strong cognitive engagement (Cavanagh & Frank, 2014; Sauseng, Tschentscher, & Biel, 2019).

Although few studies have attempted to deepen the understanding on the temporal dynamics of FMθ activity, most of them agree that its time profile could reflect different mechanisms of cognitive control and the different PFC areas involved for it (Cooper et al., 2019; Sauseng, Tschentscher, & Biel, 2019). Here, we characterize FMθ activity time profile for the planning condition as more demanding and therefore requiring a higher extent of cognitive control. As for the control condition, it was characterized as requiring cognitive control to some extent but lesser than the planning condition. The FMθ temporal dynamic activity during the planning condition was characterized as an early increase between the 4-8 Hz band that progressively grows over the first four seconds. On the other hand, the control condition showed just a transient stimulus-locked broadband increase (**Figure 4** **and** **11**).

Here, we have shown, for the first time, that as others EFs, FMθ also emerges during cognitive planning, and its temporal dynamics may be a marker of cognitive control. Additionally, source analysis confirmed that the main sources of this FMθ activity are FS and ACC (**Figure 9**–**11** and **Table 7**). The SF region is located in the superior part of the PFC and it has been described to be involved in a variety of functions associated to cognitive control functions, i.e. working memory (Boisgueheneuc et al., 2006, Owen, 2000, Owen et al., 1998, Petrides, 2000), attention (Corbetta et al., 2008, Fox et al., 2006) and sensorimotor control-related tasks (Chouinard and Paus, 2010, Martino et al., 2011, Nachev et al., 2008). Since planning requires working memory and attention, the FMθ activity observed in this region may reveal the participation of these higher order cognitive functions to support the process of planning. Furthermore, the SF region is anatomically and functionally connected to the dorsolateral PFC and the cingulate cortex through the cingulum (Beckmann et al., 2009). In this study, FMθ activity was also observed in the ACC and mACC. The dorsal part of the ACC has been associated to cognitive control, i.e. conflict monitoring (Botvinick, Cohen, & Carter, 2004; Kerns et al., 2004; Sohn et al., 2007, Ursu et al., 2009), error detection (Carter et al., 1998; Gehring & Fencsik, 2001, Pourtois et al., 2010), response selection (Turken & Swick, 1999; Awh & Gehring, 1999; Paus, 2001), and attentional control (Aarts & Roelofs, 2011; Orr & Weissman, 2009; Crottaz-Herbette & Menon, 2006; Luo et al., 2007). In the context of cognitive planning, it is required to implement these functions in order to properly elaborate a feasible plan. Altogether, we postulate that the FMθ activity from SF and ACC, by exerting working memory, attention and monitoring function, might be aiding cognitive planning by contributing to the dynamic internal elaboration of a plan. These results are in agreement with the general consensus of the existence of cognitive control core functions (Lehto et al., 2003; Miyake et al., 2000) like working memory, inhibitory control, attention, upon which higher order cognitive control functions are built such as reasoning, problem solving and cognitive planning (Collins & Koechlin, 2012; Lunt et al., 2012).

## 5. Conclusions

The present study evaluated a novel cognitive planning task with behavioral and electrophysiological measurements. Results suggest that the proposed planning task is optimal to evaluate planning, and that it induced FMθ activity originating in the PFC (SF and ACC) after 500 ms of planning period onset. We characterized both spatial and temporal dynamics of this activity during planning. It is known that planning requires cognitive control, and thus the findings in this work are in accordance with the broad body of evidence supporting the role of FMθ activity in cognitive control.

## 6. Acknowledgments

This research was financially supported by the doctoral scholarship program *Becas de Doctorado Nacional, año 2015* of CONICYT 21150295, by FONDECYT *regular* grant 1180932 and the Institut Universitaire de France (IUF).

We want to thank professor Pablo Billeke for his feedback on the paradigm design. Thanks to Josefina Ihnen, Andrea Sánchez, Ishani Thakkar, Gonzalo Boncompte, Brice Follet and Daniela Santander for their feedback.

## 8. Supplementary Material

**Table S1.** Normality Distribution Results

| Parameters* | D'Agostino & Pearson omnibus |  |
| --- | --- | --- |
|  | K2 | p-value |
| RT Planning | 4.18 | 0.12 |
| RT Execution Planning | 1.60 | 0.45 |
| Accuracy Planning | 6.20 | 0.05 |
| RT Control | 1.55 | 0.46 |
| RT Execution Control | 1.80 | 0.41 |
| Accuracy Control | 20.50 | < 0.0001**** |

RT: Reaction Time; \*Values of RT used were in seconds and values of accuracy were percentage of correct responses.

**Table S2.** Internal Consistency of tasks according to Alpha of Cronbach coefficient

| Parameters | Alpha |
| --- | --- |
| RT Planning | 0.95 |
| RT Execution Planning | 0.95 |
| RT Control | 0.97 |
| RT Execution Control | 0.97 |

RT: Reaction Time; \*Values of RT used were in seconds and values of accuracy were percentage of correct responses.

**Table S3.** Homoscedasticity Analyses

| Parameters |  | Levene's test |  |
| --- | --- | --- | --- |
|  |  | F | p-value |
| RT Planning | RT Control | 0.81 | 0.37 |
| RT Execution Planning | RT Execution Control | 1.04 | 0.31 |
| Accuracy Planning | Accuracy Control | 6.31 | 0.02 |

RT: Reaction Time; \*Values of RT used were in seconds and values of accuracy were percentage of correct responses.

**Table S4.**
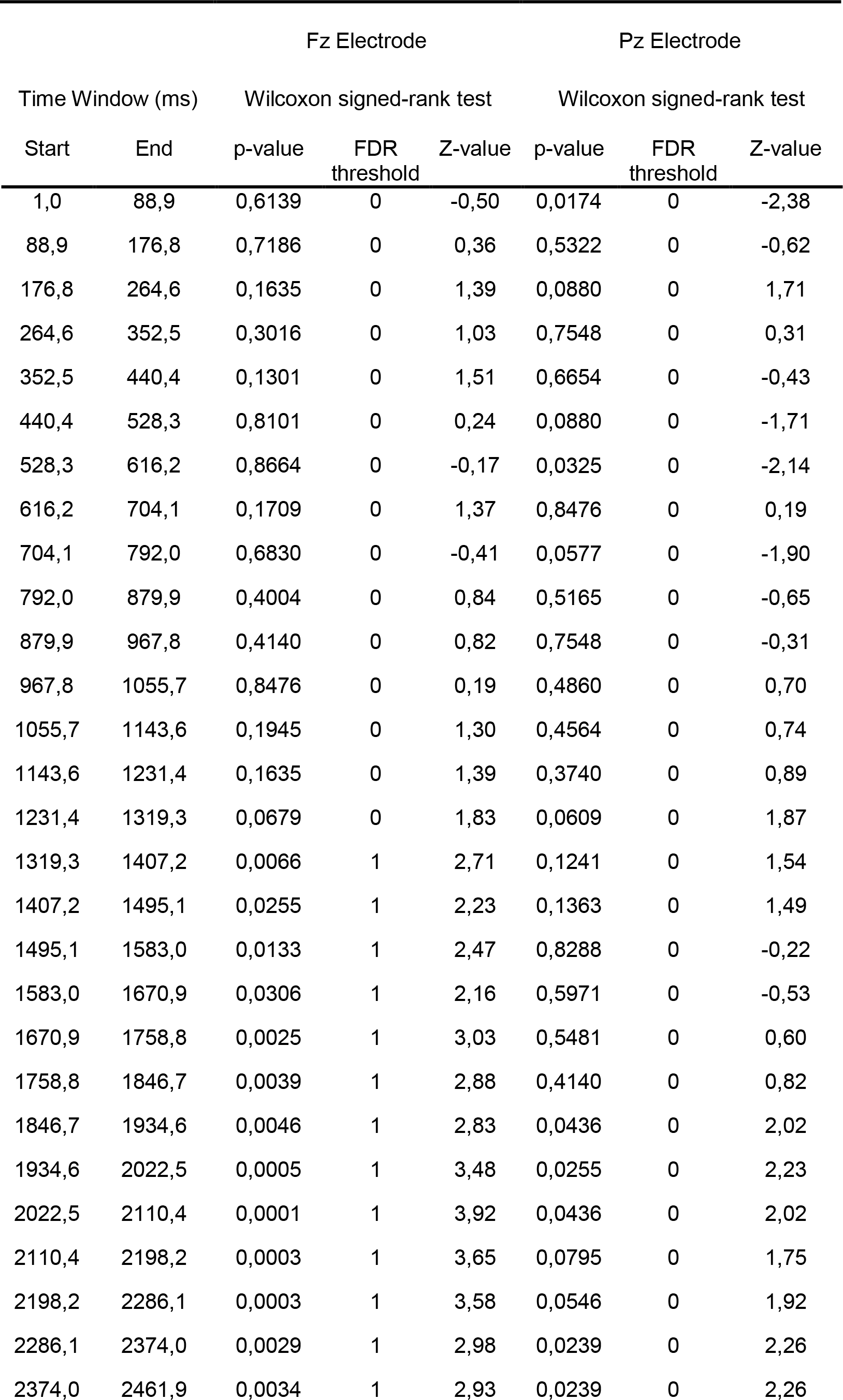
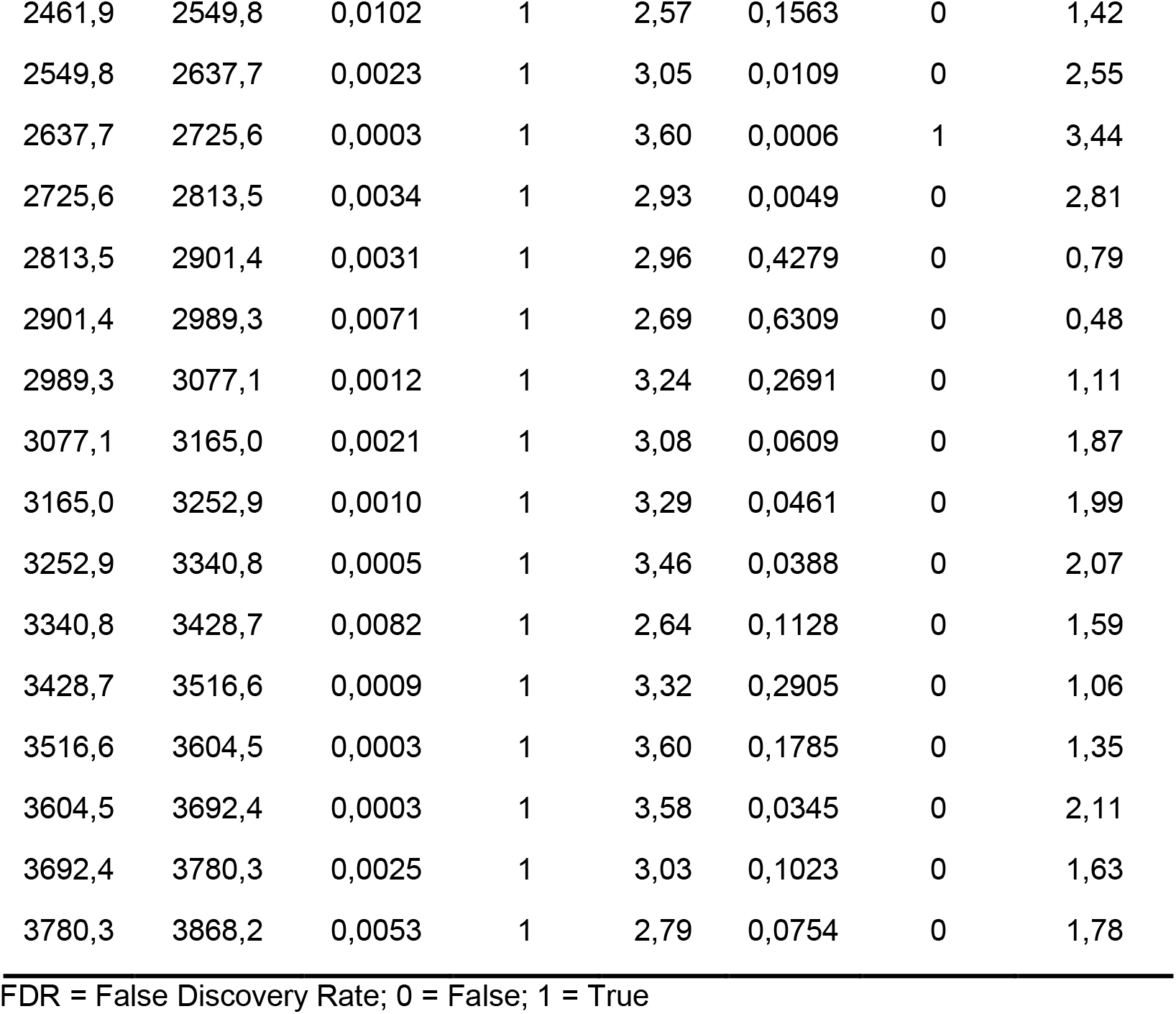
Time Slice Theta Activity: Statistics Analyses Over Time

